# Past environments modulate response to fluctuating temperatures in a marine fish species

**DOI:** 10.1101/2025.03.03.641303

**Authors:** Christelle Leung, Joëlle Guitard, Caroline Senay, Audrey Bourret, Denis Chabot, Geneviève J. Parent

**Author notes:** Correspondance: Christelle Leung, Geneviève Parent: geneviè.

## Abstract

The rise in ocean temperatures predicted due to the global warming will impact the survival and structure of various marine organisms, in particular ectothermic organisms. Phenotypic plasticity enables species to cope with environmental changes, providing a vital buffer for evolutionary changes. Yet, the dynamics and the molecular mechanisms underpinning these plastic responses remain largely unexplored. Here, we assessed the impact of acclimation environment on organisms capacity for thermal plasticity. We conducted a genome-wide transcriptomic analysis on the Acadian redfish, *S. fasciatus*, exposed to four temperatures (2.5, 5.0, 7.5 and 10.0 ℃) over a long-term period (up to 10 months) followed by an acute temperature change (24 hours), simulating natural fluctuation condition the species could encounter. Our results showed a dynamic transcriptional response to temperature involving various genes functions. The rapid response to temperature shifts, coupled with the sustained expression of specific genes over an extended period highlighted the species’ capacity for plastic response to temperature changes. We also detected a significant effect of the interaction between the long and short terms temperature exposure on gene expression, highlighting the influence of the past environment on response to acute temperature changes. Specifically, fish acclimated to higher temperatures demonstrated an increased stress-related response to environmental fluctuations, as evidenced by both the shape of their reaction norms and the implication of stress-related gene functions. This result suggests that temperature conditions predicted for the near future in the Northwest Atlantic will trigger less adaptive plasticity to environmental fluctuations, highlighting the species’ vulnerability to ocean warming.

## Introduction

Under all climatic change scenarios studied by the IPCC, temperature is projected to rise, both in term of average and intensity of fluctuations (Diffenbaugh et al., 2017; Hoegh-Guldberg et al., 2018; IPCC, 2022; Oliver et al., 2018; Seneviratne et al., 2021). This phenomenon also occurs in the oceans, driving major shifts in marine biodiversity, mainly impacting marine species distribution (Perry et al., 2005) or diminishing organisms’ performance, fitness and survival (Poloczanska et al., 2013; Ramírez et al., 2017; Smale et al., 2019; Smith et al., 2023). The adverse effect of ocean warming may be particularly severe for ectotherm organisms which lack effective physiological mechanisms to control body temperature and with limited thermal safety margin (Alfonso et al., 2021; Angilletta, 2009; Pinsky et al., 2019). Prediction of future marine communities requires therefore a better understanding of the different mechanisms underlying the organisms’ ability to respond effectively to environmental changes, a critical aspect for their persistence in natural environments.

Phenotypic plasticity is the capacity of a given genotype to develop different phenotypes according to environmental conditions (Scheiner, 1993). Adaptive plasticity reflects the development of the suitable phenotype in anticipation to future environmental conditions, allowing an increase in fitness under fluctuating and heterogeneous environments (Botero et al., 2015; West-Eberhard, 2003). Thus, plasticity can represent a powerful and effective mechanism to cope with short-term environmental changes, owing a certain predictability of the environmental changes as a key determinant of its adaptiveness (Gavrilets & Scheiner, 1993; Leung et al., 2020; Reed et al., 2010; Tufto, 2015). Understanding an organism’s capacity for plasticity needs a thorough examination of the temporal dynamics of environmental conditions. In particular, past environments can influence the initial phenotype, determining the extent of phenotypic change required to produce a more adapted phenotype in a new environment (Fey et al., 2019; Kremer et al., 2018). These past environments can also provide valuable cues for phenotype adjustment in new environments, recalibrating triggers for phenotypic changes when initial cues becomes unreliable, thereby avoiding maladaptive responses (Grieco et al., 2002). Considering past environmental conditions and the dynamics of plasticity is thus essential for understanding how organisms might cope with changing environments.

Various hypotheses have been proposed in physiology to explain how past environment influence organisms’ response to rising temperatures. For example, the Beneficial Acclimation Hypothesis (BAH) stipulates that organisms acclimatized to a specific environment would perform better or have greater fitness in that environment compared to organisms acclimated to other environments (Leroi et al., 1994). Individuals acclimated to elevated temperature over an extended period, as projected with the ongoing ocean warming scenario, would consequently be more tolerant to the environmental stress caused by the upcoming rise in temperature. Alternatively, the Optimal developmental Temperature Hypothesis (OTH) proposes that organisms acclimated to an optimal temperature will maintain higher relative fitness when confronted with a range of temperatures, compared to organisms acclimated at temperature either lower or higher than this optimum (Huey et al., 1999; Zamudio et al., 1995). These hypotheses have been the subject of several studies (e.g. Bennett & Lenski, 1997; Klepsatel et al., 2019; Steigenga & Fischer, 2009; Wilson & Franklin, 2002). However none have tested the effect of past environment on organisms’ capacity for plasticity. In other words, it remains unclear whether individuals acclimated to different temperatures would display varying levels of plasticity when coping with the same temperature challenges.

For this purpose, we studied the thermal plasticity of the Acadian redfish, *Sebastes fasciatus*. This species from the *Sebastes* genus generally inhabits deep waters (250-500 m) of the temperate Northwest Atlantic Ocean (Gascon, 2003). This commercial fish is exposed to warming environmental conditions. In the Estuary and Gulf of St. Lawrence (hereafter St. Lawrence System), the average of bottom temperature maximum has increased from 5.2 °C in 2009 to 6.9 °C in 2023 (Galbraith et al., 2024). During the last decade, shifts in species distributions, changes in ecosystem productivity and an increase of *Sebastes* species biomass were observed both in the St. Lawrence System (from 100,000 to 4,300,000 tonnes, Senay et al., 2023) and other fisheries management areas of the Northwest Atlantic (DFO, 2023). The factors contributing to the increase in biomass of *Sebastes* spp. remain unclear, but the rising temperature has been identified as a potential driving force, particularly affecting the early developmental stages within the St. Lawrence System (Burns et al., 2020, 2021). This species has to deal with acute temperature challenges, both in its development stage and as an adult. During development, while migrating to the seabed to continue its growth, juvenile *S. fasciatus* cross a cold intermediate layer at depths of 50 to 120 meters, characterized by temperatures around 0–1 °C (Galbraith et al., 2024; Senay et al., 2023). As adult, *S. fasciatus* perform daily migrations for feeding, during which they could be exposed to cooler temperature if approaching the cold intermediate layer at night (Gascon, 2003). Consequently, this species is likely to experience temperature fluctuations due to these behaviors. How past acclimation temperatures, whether during the larval stage or as an adult, influence the species’ ability to face acute temperature changes remains unclear.

In this study, we investigated the capacity for plasticity of an ectotherm species to cope with rapid thermal changes when acclimated to different temperatures. Plasticity applies for many phenotypic traits from molecular level to more integrated phenotypes encompassing physiological mechanisms, morphology, and behavior. Here, we assessed the level of plasticity of the most fundamental level at which the genotype gives rise to the phenotype, that is gene expression, under warming ocean scenarios. Since we are using a non-model species, we first characterized the molecular mechanisms underpinning the thermal plasticity of *S. fasciatus*. Then, we assessed the effect of two different acclimation temperature on short-term (24 h) temperature exposure. While long-term temperature exposures mimic current (Galbraith et al., 2024) and predicted temperatures (Lavoie et al., 2020), the short term temperature changes simulate conditions that could be experienced by redfish during its daily migration for feeding (Gascon, 2003). Functional analysis of differentially expressed transcript also allowed to shed light on the genes functions involved in *S. fasciatus* response to temperature variation.

## Materials and Methods

### Experimental design

Tissue samples were collected from *S. fasciatus* in a tank experiment described in Guitard et al. (2025). Briefly, fish were captured near *les Escoumins*, Québec, Canada (48.317801°N, 69.413287°W) in fall 2019. SCUBA divers caught *ca.* 16–19 cm fish with dip nets at a depth of 25–30 m. Fish were placed into cages (52 x 31 x 31 cm), that could hold up to 30 individuals, and full cages were left at a depth of 10–15 m depending on the tide for at least 12 h, but up to 96 h, to reduce barotrauma. Fish were then transported in oxygenated tanks to the Maurice Lamontagne Institute in Mont-Joli, Québec, Canada.

The experimental set-up was composed of eight circular tanks (1 m diameter; 760 L) arranged into four temperature lines (2.5, 5.0, 7.5, 10.0 ℃) with pH maintained at 7.75 (total scale). The four temperatures were selected based on the typical living conditions of *S. fasciatus*, which is around 5 °C. The experimental temperature range included both colder and warmer conditions, with the warmest conditions reflecting the 10 °C expected for 2061 according to modelling scenarios for the St. Lawrence System (Lavoie et al., 2020). A total of 25 fish were randomly distributed in each tank. Fish were acclimated to the experimental tanks for a week at the same rearing conditions (5 ℃). Then, temperatures were adjusted in increments of 0.5 ℃ per day until all experimental temperatures were reached (*e.g.,* 2.5 weeks for 10.0 ℃). Experimental conditions were reached on January 25^th^, 2021, and the end of the experiment was scheduled on June 29^th^, 2021. However, slow growth rates and many mass losses were observed during this five month experiment in all treatments, due to a feeding protocol that did not result in *ad libitum* feeding and possibly a suboptimal fish density (see Guitard et al., 2025). Thus, more fish were added to each experimental tank after a gradual temperature adjustment as described previously. The additional fish reached the experimental temperature on July 16^th^ and the growth experiment ended on October 20^th^. This adjustment in the protocol resulted in fish undergoing long-term acclimation periods of either 10 or 3 months. Each fish was uniquely identified using a Passive Integrated Transponder (PIT tag).

To assess the short-term transcriptional response to temperature change, fish from 5.0 and 7.5 ℃ treatment were subjected to ± 2.5 ℃ temperature changes for 24 h. The day before the end of the experiment, floating basket were installed in each experimental tank. For fish acclimated to 5.0 ℃ and 7.5 ℃, nine fish were randomly selected in each tank. Three fish stayed in the basket of the same tank, three fish were transferred to 2.5 ℃ colder temperature (*i.e.* 2.5 and 5.0 ℃ tank for fish from 5.0 and 7.5 ℃, respectively), and three fish were transferred to 2.5 ℃ warmer temperature (*i.e.* 7.5 and 10.0 ℃ for fish from 5.0 and 7.5 ℃, respectively). Three fish were randomly selected in the 2.5 and 10.0 ℃ treatments and placed in the basket of the same tank. This last step ensured that all fish were subjected to the same basket transfer manipulation. After 24 h, fish were taken one by one from the floating basket and euthanized by blunt force trauma followed by exsanguination. A total of 48 fish were sacrificed, which included 24 from the long-term exposure experiment (4 temperatures × 2 tanks × 3 fish) and 24 from the short-term exposure experiment (2 acclimation temperatures × 2 acute temperatures × 2 tanks × 3 fish). For each individual, muscle tissue was sampled and flash frozen in liquid nitrogen before being stored in −80 °C until nucleic acid extraction. Genetic analysis confirmed that all individuals were *S. fasciatus* (Fig. S1A), and mostly belong to a single genetic group (Fig. S1B; see Supplementary Material 1 for more details).

Fish collection in their natural habitat was done under the Canada parks permit (SAGMP-2019-33741). Experimental methods complied with the regulations of the Canadian Council on Animal Care and were approved by the Maurice Lamontagne Institute animal care committee (Certificates 19-6B, 19-6C and 19-7B).

### RNA extraction, sequencing and bioinformatic pre-processing

RNA extraction of the 48 redfish was carried out on *ca.* 2 g of white muscle tissue, using Qiazol RNA extraction reagent (Qiagen) following the manufacturer’s protocol. RNA-sequencing (seq) was used to investigate the changes in the entire transcriptome. Library construction, using NEB mRNA stranded library preparation kit, and high-throughput sequencing steps were performed by Génome Québec (Montreal, QC, Canada). Two sequencing batches were performed using Illlumina NovaSeq 6000 S4 PE100 and PE150, for respectively 33 and 21 libraries including six samples in common for both sequencing batches. No sequencing batch effect on the measured transcript expression levels was detected (Fig. S2).

The RNA-seq raw reads quality was assessed using FastQC version 0.11.9 (Andrews, 2010) and MultiQC version 1.9 (Ewels et al., 2016). They were then trimmed with Trim Galore! version 0.6.6 (Krueger, 2015) to remove adaptor sequences and to obtain a minimum Phred score of 30 and a minimum length of 20 base pairs (bp). We aligned the trimmed reads on a *Sebastes fasciatus* genome (GeneBank accession: JBJQUQ000000000) using the splice-aware alignment program HISAT2 version 2.2.1 (Kim et al., 2015), with default parameters for paired-end reads.

*De novo* transcriptome reconstruction was performed with Stringtie version 2.2.1 (Pertea et al., 2015) based on the aligned reads for all the libraries and in a conservative mode. This procedure ensured complete transcripts identification from the experimental samples. Structural annotation of the genome was performed by using the Stringtie transcript merge usage mode to assemble transcripts from the obtained *de novo* transcriptome reconstruction of each library. This procedure resulted in the generation of a unified and non-redundant set of isoforms across the RNA-seq samples. We, then, quantified the number of reads per transcript with FeatureCounts version 2.0.3 (Liao et al., 2014) using the aligned reads from HISAT2 and the transcript structural annotation file from Stringtie transcript merge usage mode.

Gene Ontology (GO) assignments were used to classify the functions of *S. faciatus* transcripts. The transcriptome functional annotation was thus obtained based on sequence homology to known sequence database. NCBI’s *blastx* tools version 2.12.0+ (Camacho et al., 2009) with an e-value cut-off of 0.001 was used to compare the transcript sequences against the SwissProt protein database (downloaded May 30^th^, 2022). The UniProt information, including the associated GO terms corresponding to the hit, was then retrieved from the UniProt website.

### Statistical analyses

Gene expression variation partitioning and differential expression analyses were performed using the R package *vegan* version 2.6-4 (Oksanen et al., 2022) and the Bioconductor’s package *DESeq2* version 1.40.2 (Love et al., 2014). We quantified the proportion of the total gene expression variation explained by long-term and acute temperature exposures by applying partial redundancy analyses (pRDA; Borcard et al., 1992). Samples were first filtered according to the duration of the exposition to a given temperature: long-term or acute exposure conditions encompassed individuals exposed to the four temperatures for more than three months or 24 h, respectively. We then used the normalized transcript count matrix (*i.e.*, applying a variance stabilizing transformation (VST) to the number of reads per transcript matrix, as implemented in *DESeq2*) as the response variables and sampling temperatures as the explanatory variables. In particular, as fish could have been exposed to 3 or 10 months for the long-term exposure conditions (see experimental design described above), the effect of different durations in a given temperature was also tested through a pRDA. For short term exposure conditions, we used as explanatory variables the acclimation and sampling temperatures, and their interaction since fish were acclimated at 5 or 7.5 °C before been transferred at ± 2.5 °C for 24 h.

We identified transcripts that showed changes in expression between temperatures by building a general linear model as implemented in *DESeq2* and using the Wald test (Love et al., 2014). Transcripts with FDR < 0.05 (*P*-value after Benjamini-Hochberg adjustment) and |log2FC| > 1 were considered as differentially expressed. We used fish from 5.0 °C exposure as the control temperature for the long-term exposure experiment, as they have been acclimatized at this temperature from their arrival to the laboratory facilities until the experimental treatments. This procedure allowed us to test for differentially expressed transcripts (DETs) when temperature is reduced (from 5.0 °C to 2.5 °C) or increased (from 5.0 °C to 7.5 °C or 10.0 °C). Similarly, for acute temperature exposure treatment, redfish acclimatized at 5.0 °C or 7.5 °C were used as control conditions to test for DETs between temperatures of both control conditions and sampling temperatures. This procedure allowed us to identify transcripts involved in reduced (−2.5 °C) or increased (+ 2.5 °C) temperature within 24 h. DETs with similar expression pattern on the basis of Kendall rank correlations (at least 15 transcript per cluster, correlation > 0.7, testing the hypothesis of correlation for a confidence interval of 95%) were then grouped together, based on the normalized count matrix using the *DEGreport* version 1.36.0 package (Pantano, 2023).

We finally used the Gene Ontology (GO) assignments to classify the functions of *S. fasciatus* identified as DETs across temperatures. Enriched GO terms of the DETs were assessed using the ‘classic’ algorithm from the *topGO* version 2.52.0 package (Alexa & Rahnenfuhrer, 2023) and based on *P*-value generated using Fisher’s exact test method, with a threshold of FDR ≤ 0.001 after Benjamini-Hochberg adjustment, for the three GO categories (*i.e.*, biological process, molecular function and cellular component). To facilitate the interpretation of the obtained large list of GO terms associated to DETs, similar terms displaying the most probable gene function candidates were grouped based on their semantic similarity. The obtained reduced list of GO terms was then visualized using a treemap (space-filling visualization) of grouped terms using *rrvgo* version 1.12.0 package (Sayols, 2023), where the GO terms function names were retrieved from the Bioconductor genome wide annotation for Zebrafish (Carlson, 2019). Through this procedure, we could visualize the importance of a given function, as the space used by a given term is proportional to the −log10 (FDR) (*i.e.* the largest is the space, the most probable is the gene function candidate). Finally, heat-shock proteins (HSPs), that are molecular chaperones that aid in protein folding and degradation, represent a widely conserved cellular stress response mechanism to temperature changes, and play an ecological and evolutionary important role in thermal adaptation (Feder & Hofmann, 1999; Logan et al., 2015; Tomanek, 2010). We thus specifically searched for transcript annotated to HSP functions and assessed how temperature affected their expression in *S. fasciatus*.

## Results

### *De novo* transcriptome reconstruction

We obtained a total of 2.26 × 10^9^ raw paired reads, from which 98.2% remained after the trimming step. The genome structural annotation was obtained based on the trimmed reads aligned on a *S. fasciatus* assembly, with an average overall alignment rate of 97.9% (*s.d.* 0.6%) per library. The *de novo* transcriptome reconstruction resulted in 35,483 putative transcript isoforms that could be grouped into 17,772 genes present across all samples, with a median length of 8,619 bp. A total of 12,392 (69.7%) genes were characterized by a single transcript per gene. A total of 26,647 *S. fasciatus* transcripts (*ca.* 75.1% of 35,483 transcripts) were successfully annotated to a specific function based on the sequence homology to SwissProt protein database and assigned to at least one GO term and categorized into a total of 14,285 GO terms.

### Transcriptional response to temperature variation

#### Quality check and sex effect on gene expression

FastQC quality check indicated that two libraries displayed high proportion of overrepresented sequences (Fig. S3). A manual *blast* of these overrepresented sequences revealed that these two libraries displayed ribosomal RNA (rRNA) sequence contamination. This likely resulted from an incomplete or a failure of the rRNA depletion step during library preparation. These two libraries, including one duplicated sample (*i.e.* sequenced twice, once in each of the two batches), were removed for the subsequent analyses. In addition, only one randomly chosen of the remaining five duplicated samples were retained. Transcriptional responses to temperature variation were therefore performed on 47 libraries.

Due to their small size, sex of each individual were therefore genetically assessed (Fig. S4A). No sex (*adjusted R^2^* = 0.00 %; *P* = 0.669) or sex × temperature interaction (*adjusted R^2^*= 0.00 %; *P* = 0.453) effect on transcript abundances were detected with a redundancy analysis (Fig. S4), indicating no sex-specific response to temperature change. Therefore, sex was not used as covariable in the subsequent analyses.

#### Response to long-term temperature exposure

The four temperatures tested for long-term exposure had an effect on transcript levels (*adjusted R^2^* = 22.67%, *P* = 0.001; Fig. 1A). Fish exposed at 2.5 and 5.0 °C expressed similar transcript pattern while fish exposed at 10.0 °C displayed the most different transcript pattern from those at 2.5 and 5.0 °C. Fish at 7.5 °C displayed intermediate transcript profiles to those of 2.5/5.0 °C and 10 °C (Fig. 1A). Pairwise comparison analysis of DETs confirmed these results, as the smallest number of DETs was observed between 2.5 and 5 °C (n = 75 DETs; Fig. 1B) which contrasted considerably with the number of DETs between 5.0 °C and 7.5 °C (n = 733 DETs; Fig. 1B), and 5.0 and 10.0 °C (n = 3,343 DETs; Fig. 1B). Note that fish exposed for 3 or 10 months displayed no significant differences in transcript levels (*adjusted R^2^* = 1.05%, *P* = 0.102), and both conditions were thus considered as long-term acclimation.

**Fig. 1.**
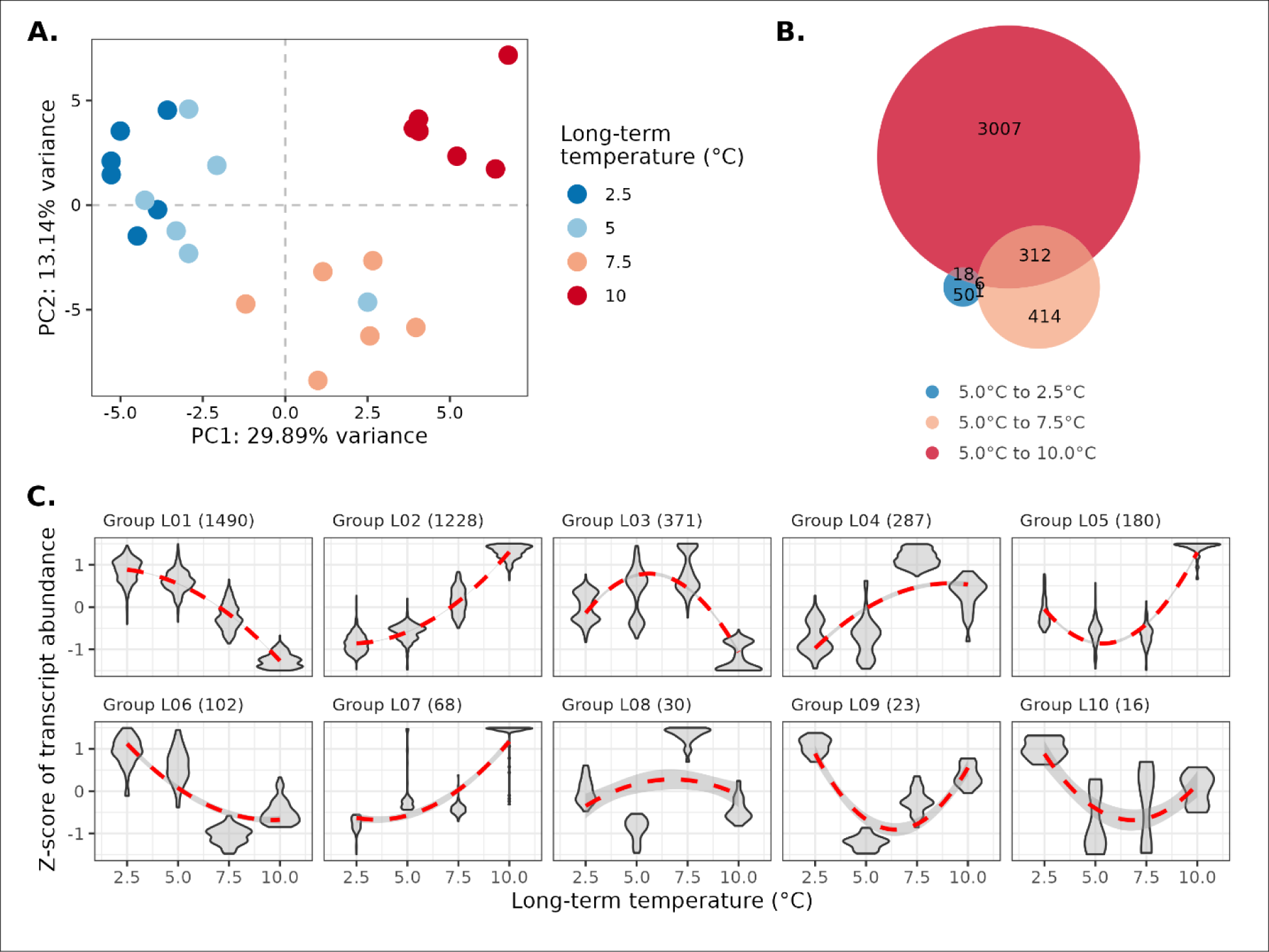
Transcriptional response to long-term temperature exposure. (A) Variation in transcript expression levels. Principal Component Analysis of transcript expression levels of individuals with long-term exposure (3 to 10 months) to different temperatures (colors). (B) Number of Differentially Expressed Transcripts (DETs). Venn diagram displays the count of DETs between the control temperature (5.0 °C) and the three other long-term temperature exposures, 2.5 °C (blue), 7.5 °C (orange) and 10.0 °C (red). (C) Groups of co-expressed transcripts for DETs among long-term temperature exposures. Numbers in parentheses indicate the number of transcripts belonging to each co-expression module.

DETs among temperatures for long-term exposure were grouped into 10 groups of co-expressed transcripts. More than 71% of these DETs displayed an almost linear effect, either decreasing (Group L01, n = 1,490 DETs, 39.26%; Fig. 1C) or increasing (Group L02, n = 1,228 DETs, 32.36%; Fig. 1C) transcript expression levels as a function of temperature. For these two groups of DETs, transcript expression levels were again more similar for fish exposed to 2.5 °C and 5.0 °C, compared to fish at higher temperature exposures. Another similar expression pattern was observed for the DETs groups L04 (n = 287, 7.56%) and L06 (n = 102, 2.69%), where a step increase or decrease in transcript expression level was detected at 7.5 °C. This result illustrated that a temperature decrease from 5.0 °C to 2.5 °C involved less gene expression changes than a temperature increase from 5.0 °C to 7.5. The expression pattern observed in DETs between temperatures from groups L03 (n = 371), L05 (n = 180), L09 (n = 23) and L10 (n = 16), totalizing 15.55% of DETs among long-term temperature exposures, was characterized by a U- or bell-shaped expression pattern where respectively higher or lower expression level was observed for fish from extreme temperatures (*i.e.* 2.5 °C and 10.0 °C) compared to those from intermediate ones (*i.e.* 5.0 °C and 7.5 °C).

#### Response to short-term temperature exposure

Transcript expression levels were also affected by the short-term temperature exposure. Variation partitioning analysis revealed that both *Acclimation* temperature (*i.e.* where fish spent up to 10 months; *adjusted R^2^* = 3.43%, *P* = 0.005) and *Sampling* temperature (where fish spent 24 h after the acclimation step, *adjusted R^2^* = 2.49%, *P* = 0.008) affected transcript abundance (Fig. 2). This result confirmed the thermal plasticity of *S. fasciatus* through transcript regulation. The response to temperature changes was asymmetrical as a function of the direction of temperature changes. For fish from both acclimation conditions (5.0 and 7.5 °C), a greater number of DETs were detected in fish subjected to a temperature increase of 2.5 °C compared to those exposed to a temperature decrease of 2.5 °C (Fig. 2B). Interestingly, we also observed that changes in expression levels of certain genes, which occurred during the first 24 h, were maintained over a period of 10 months (e.g. Groups S01, S04 and S08 at 5.0 °C, and S02, S05 and S07 at 7.5 °C, Fig. 2C).

**Fig. 2.**
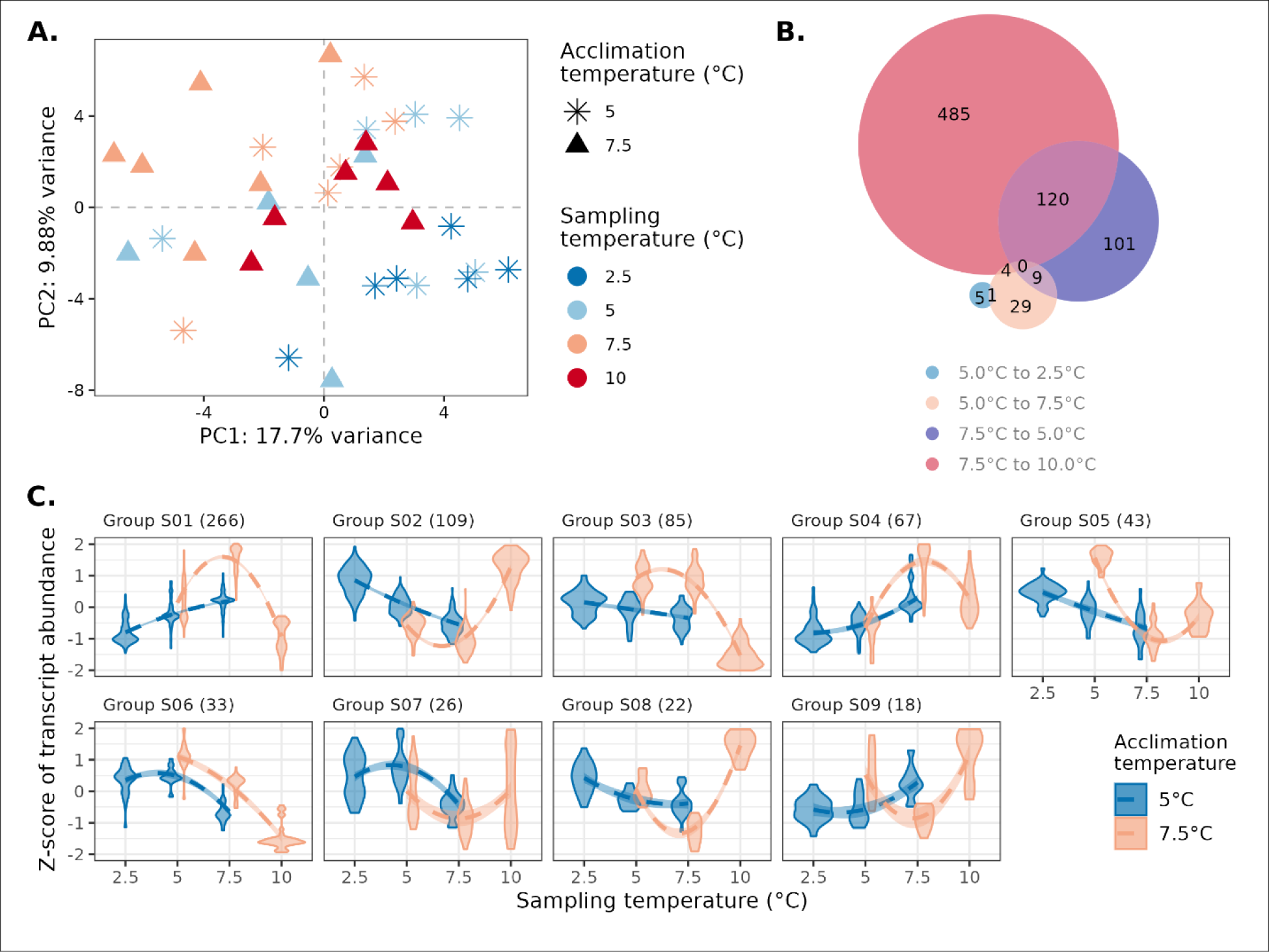
Transcriptional response to short-term temperature exposure. (A) Variation in transcript expression levels. Principal Component Analysis of transcript expression levels of individuals acclimated or long-term exposed to (shape) and sampled or short term exposed to (sampling, color) different temperatures. Different acclimation and sampling temperature indicates that individuals were subjected to a short-term temperature change. (B) Number of Differentially Expressed Transcripts (DETs). Venn diagram displays the count of DETs between the acclimation temperature (5.0 or 7.5 °C) and sampling temperature (2.5 (blue) and 7.5 °C (orange) for fish acclimated at 5.0 °C, and 5.0 (purple) or 10.0 °C (red) for fish acclimated at 7.5 °C). (C) Groups of co-expressed transcripts for the total DETs obtained in (B), represented according to sampling temperatures and grouped by acclimation conditions (colors). Numbers in parentheses indicate the number of transcripts belonging to each co-expression module.

The *Acclimation* × *Sampling* temperatures interaction also explained a significant part of the transcript abundance variation (*adjusted R^2^* = 2.49%, *P* = 0.020). This suggests that the short-term response to temperature changes depend on long-term acclimation temperatures. The effect of acclimation in the short-term response is evidence by the significantly fewer DETs among sampling temperature for fish acclimated at 5.0 °C (n = 48 DETs) compared to those acclimated at 7.5 °C (n = 719 DETs; Fig. 2B). This observation was also made at the level of transcript reaction norms (Fig. 2C). For DETs associated with the short-term specific response, fish acclimated at 5.0 °C showed relatively linear reaction norms (Fig. 2C, blue lines). In contrast, the majority of reaction norms for fish acclimated at 7.5 °C were characterized with bell or U-shaped patterns. This indicates for a similar gene expression response to either a 2.5 °C increase or decrease in temperature (all except S06; Fig. 2C).

### Functional analysis of DETs

We used a Gene Ontology (GO) enrichment analysis to examine whether the transcriptional plasticity in response to long- and short-term temperature changes in *S. fasciatus* involved different functional requirements. We investigated the function of different groups of DETs: those exclusive to long-term response (n = 3,501 DETs), those common to both long- and short-term response to temperature changes (n = 307), and those exclusive to short-term response. However, for the short-term response, we further differentiated between DETs detected in samples acclimated at 5.0 °C (n = 31) and 7.5 °C (n = 428) (Fig. S5), which allowed us to better assess the gene function associated to the significant *Acclimation* × *Sampling* temperature interaction.

DETs associated to exclusively long-term response to temperature changes involved mainly regulation of muscle activity including the regulation of muscle contraction, muscle tissue morphogenesis and cell physiological processes like ion transport, cGMP biosynthesis or amino-acid transport (Fig. 3). We also performed a GO enrichment analysis on groups of co-expressed that displayed a down- (group L01; Fig. 1C) and up- (group L02; Fig. 1C) regulation of gene expression as temperature increased. Genes associated to transmembrane transport, muscle contraction, oxidoreductase and hormone activities were downregulated at warmer temperatures (Fig. S6). At the opposite, genes involved in cell growth and organization, protein folding or biogenesis were upregulated at higher temperature (Fig. S7). Interestingly, we also observed higher expression of transcripts associated to histone H3R17 methyltransferase activity (Fig. S7B) at higher temperatures, suggesting fine-tuning gene expression regulation *via* epigenetic process.

**Fig. 3.**
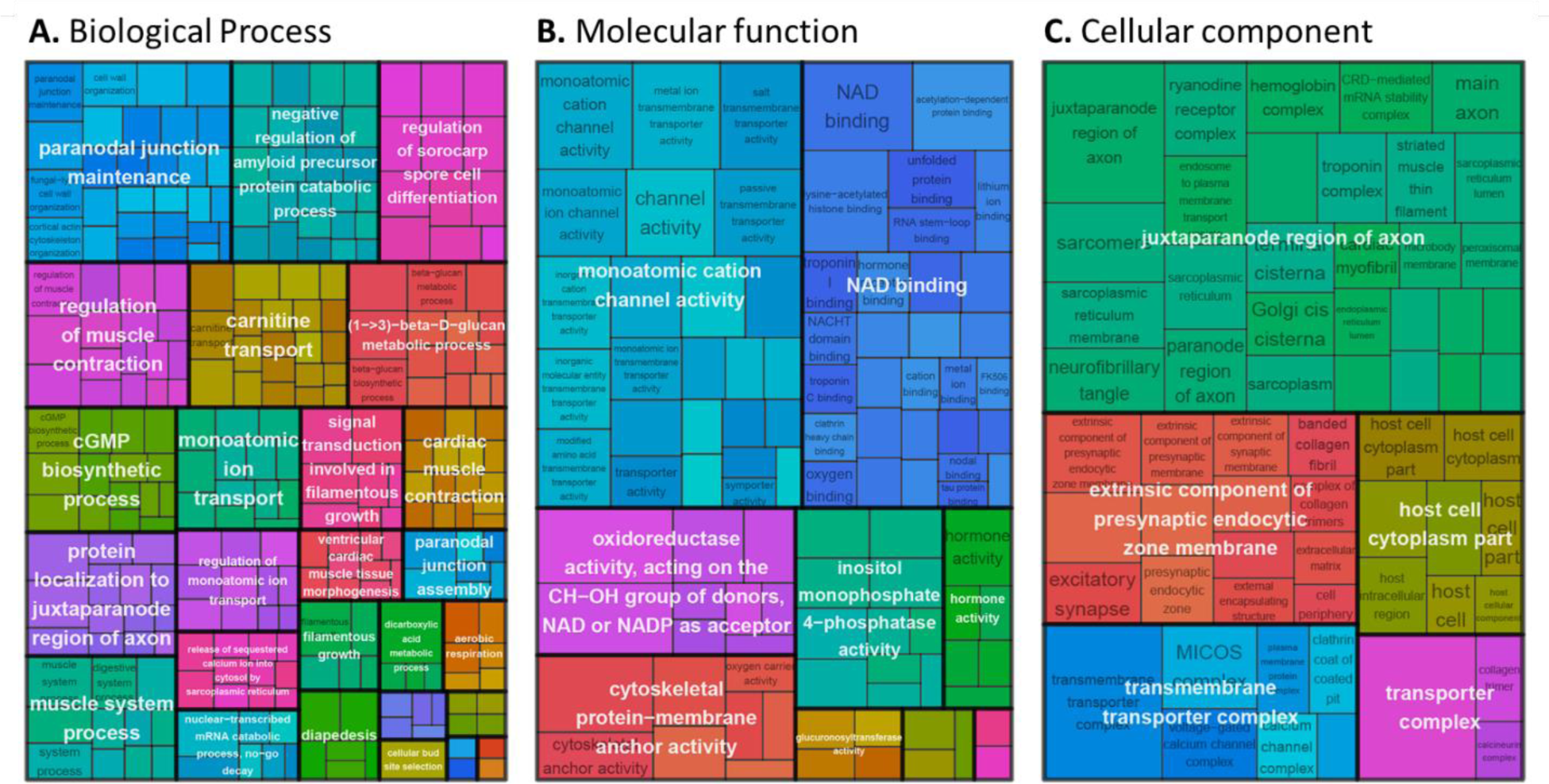
GO terms associated to exclusively long-term response to temperature change. Treemap for (A) biological process, (B) molecular function and (C) cellular component where GO terms were grouped (color) based on their semantic similarity, and the space used by the term is proportional to the −log10(adjusted *P-value*), hence the gene function candidate probability.

DETs in common between long- and short-term response to temperature changes involved genes associated to stress response. This included mainly gene function linked to stress-induced cellular senescence and cell communication (Fig. 4). Surprisingly, even if the tissue used in the current study was white muscle, we still observed a GO enrichment for the optic cup formation for eyes development (Fig. 4A). This represented a group of 143 DETs, distributed among 26 reduced GO terms. These DETs were involved mostly in the long-term and short-term temperature response for sample acclimated at 7.5 °C, as it represented 116 and 11 DETs, respectively. But it was also a common response between the long- and short-term temperature response for fish acclimatized at 7.5 °C, with 15 common DETs between both conditions, while short-term temperature response for fish acclimated at 5.0 °C only involve one DET associated to this eye development biological process.

**Fig. 4.**
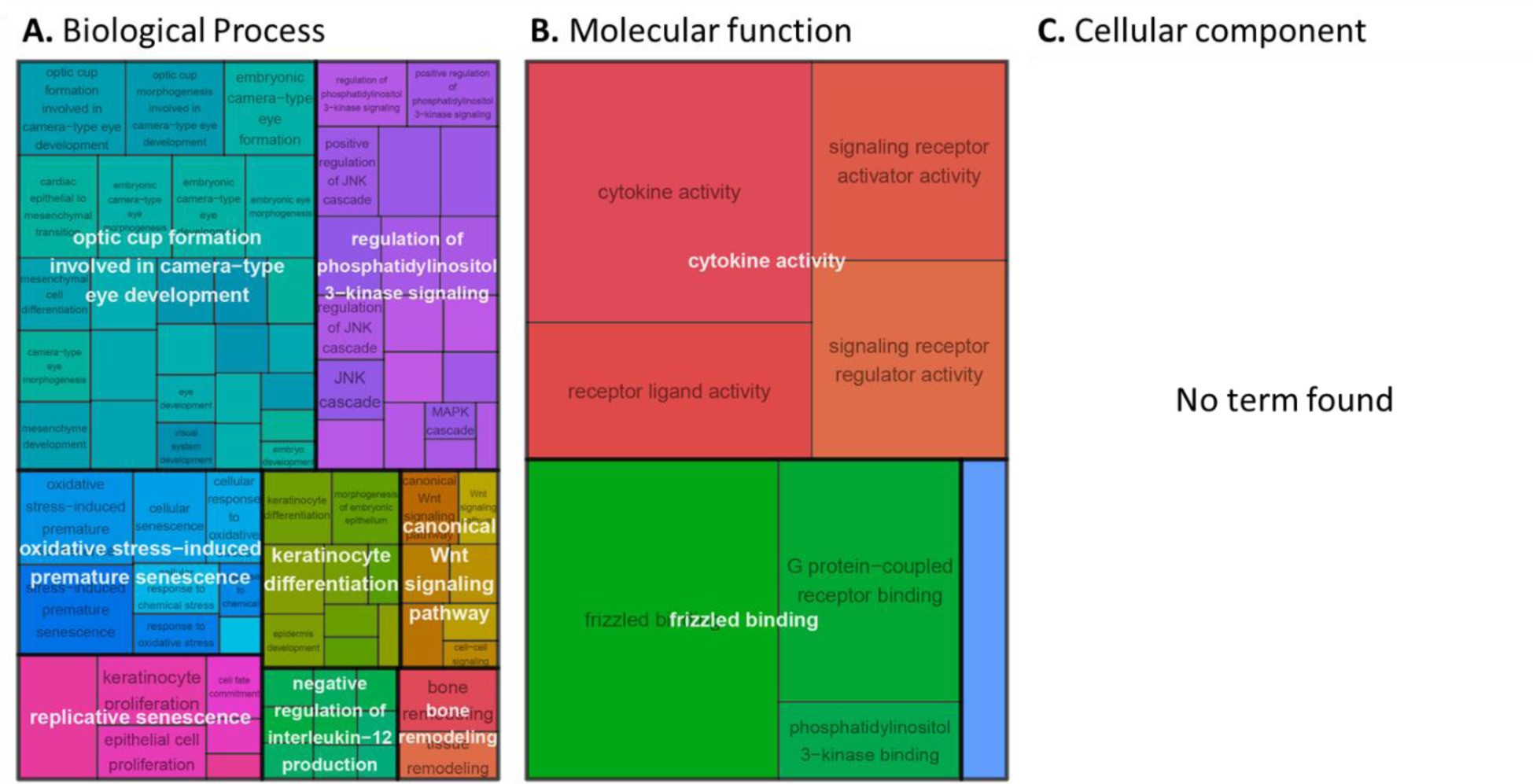
GO terms associated to DETs in common between long and short term response to temperature change. Treemap for (A) biological process, (B) molecular function and (C) cellular component where GO terms were grouped (color) based on their semantic similarity, and the space used by the term is proportional to the −log10(adjusted *P-value*), hence the gene function candidate probability.

Function of genes exclusively expressed in response to short-term temperature changes was detected to be acclimation temperature dependent. Samples acclimatized at 5.0 °C involved DETs that were associated with energy production (e.g. glucose import, hormone secretion, dATP metabolic process, Fig. 5A), and gene expression regulation (e.g. histone acetyltransferase (Gong et al., 2020), Fig. 5B). Samples acclimatized at 7.5 °C also expressed genes linked to muscular cell and gene regulation. However, they also mainly displayed stress-related gene expression, with DETs associated with cell signaling, immune cells communication and cell killing regulation (Fig. 5D to F). Specifically, the search of the key word “stress” in GO term enrichment results revealed a total of eight GO terms associated to response to temperature fluctuation for fish acclimated to 7.5 °C, while no GO terms with the keyword “stress” were detected for those acclimated to 5 °C.

**Fig. 5.**
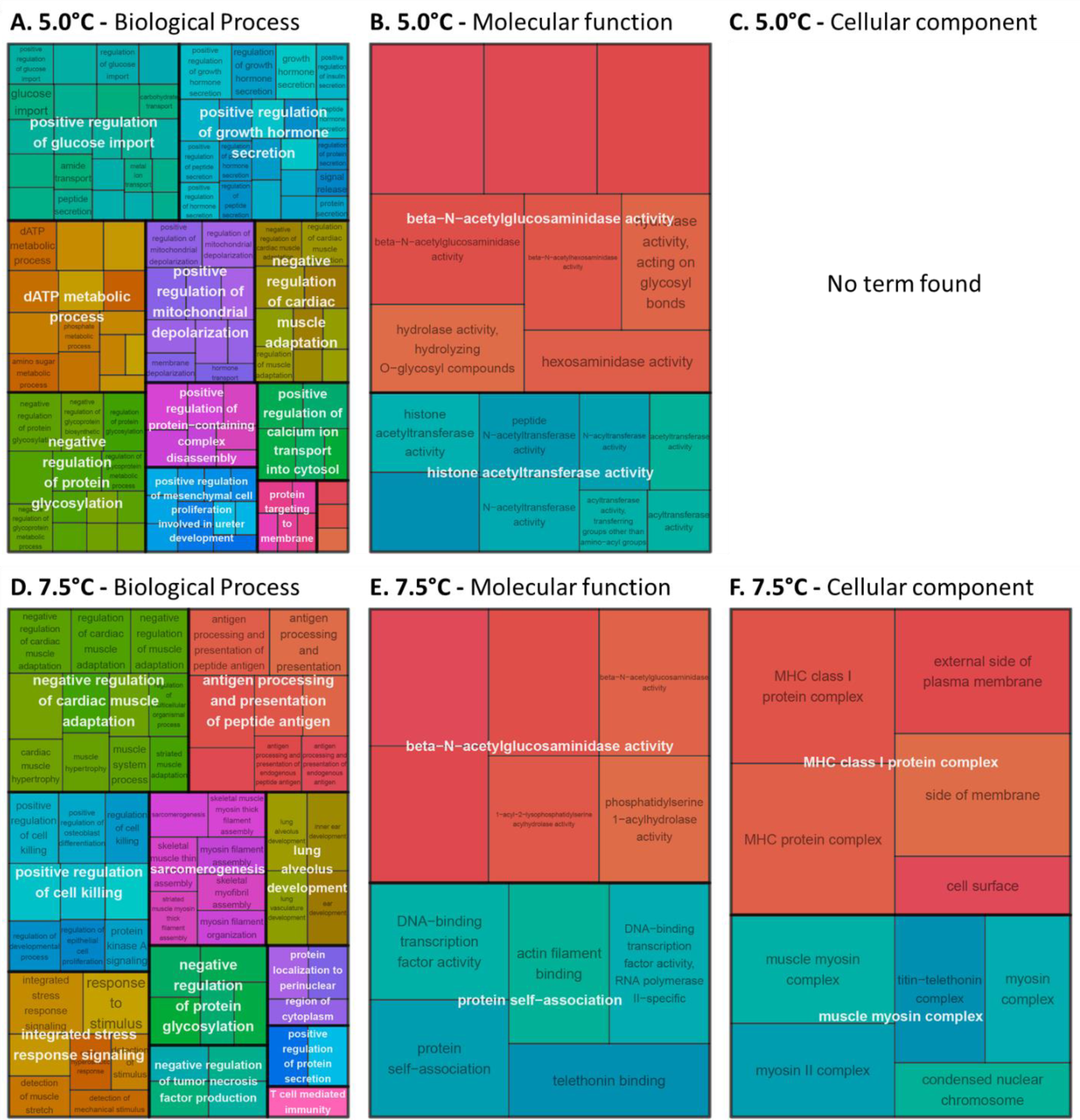
GO terms associated to exclusively short-term response to temperature changes, for fish acclimated at 5.0 °C (A, B and C) and 7.5 °C (D, E and F). Treemap for (A and D) biological process, (B and E) molecular function and (C and D) cellular component where GO terms were grouped (color) based on their semantic similarity, and the space used by the term is proportional to the −log10(adjusted *P-value*), hence the gene function candidate probability.

A targeted search for the activity of heat-shock proteins (HSPs) revealed that this gene family is indeed present and expressed in *S. fasciatus* but exhibits a weak response to temperature stress. We identified a total of 125 transcripts associated with HSPs. Of these, only 25 were found to be differentially expressed in response to long-term temperature exposure, while the majority (100 transcripts) maintained uniform expression level across all four experimental temperatures (Fig. S8).

## Discussion

In this study, we described the transcriptional plasticity of the redfish *S. fasciatus*, in response to both short- and long-term temperature changes. Plasticity in gene expression allowed the species to cope with temperatures exceeding those typically encountered in their adult habitat within the St. Lawrence System and close to the maximum temperature where the species as been observed in the wild (Scott, 1982). Gene expression changed rapidly and was maintained for short- and long-term temperature changes, respectively, highlighting the species’ capacity for a plastic response to temperature changes. Multiple gene functions were implicated in both short- and long-term responses. The acclimation temperature also influenced the response to acute temperature change. Fish acclimated to warmer temperatures displayed more short-term transcriptomic changes and associated to stress-related gene functions, in response to temperature change compared to those acclimated at colder temperatures. Our results show that capacity for plasticity does not shield this species from the impacts of climate change. On the contrary, our comprehensive understanding of the mechanistic response to temperature stress informs us about the vulnerability of this species to ocean warming.

### The plasticity of gene expression responses to short- and long-term temperature exposures

Reaction norms, which describe the range of trait changes across environmental conditions, are effective tools for assessing an organism’s capacity for plasticity. In this study, transcript expression patterns were distinct for fish acclimatized to four temperatures, illustrating the capacity for transcriptional thermal plasticity in *S. fasciatus*. The near-linear expression profile relative to temperature, encountered for most transcripts during the long-term exposure, suggests that the species copes progressively with increasing temperatures. This result aligns with previous studies showing that the majority of fish species have such capacity for plasticity following a change in environmental conditions (reviewed in Oomen & Hutchings, 2017).

Transcript profiles were more similar for 2.5 and 5.0 °C, while more different between 5.0 °C and higher temperatures (7.5 and 10 °C). This result could be an indicator of the species’ preferred temperature around 2.5 and 5.0 °C. These findings are consistent with a study that actually demonstrated the species’ best performance for growth for temperatures at 2.5 and 5.0 °C, which facilitate a manageable metabolic demand (Guitard et al., 2025). These results also reflect the temperature conditions of the last decade in the St. Lawrence system (Galbraith et al., 2024), where *S. fasciatus* encounters these specific temperature challenges (Senay et al., 2023).

The assessment of temporal dynamics of plasticity, also known as the rate of plasticity, is crucial to enhance our understanding of its adaptiveness relative to environmental changes (Burton et al., 2022; Dupont et al., 2024; Einum & Burton, 2023). Our results from the short-term exposure revealed that *S. fasciatus* responded to temperature variations within a 24-hour timeframe, highlighting this species’ ability to modulate rapidly gene expression. In addition, the levels of gene expression needed to cope with a specific temperature were maintained over time. This was evidenced by the comparable expression levels of various genes between the long-term and short-term exposures to 5.0 or 7.5 °C, the linear reaction norm after an acclimation at 5.0 °C initiated within 24 h after temperature changes and maintained over 10 months, and the lack of significant differences in gene expression patterns between samples exposed to a given temperature for 3 or 10 months. Together, these results highlight the ability of the species to rapidly fine-tune and maintain differences in gene expression in response to temperature fluctuations. Such capacity could be adaptive in fluctuating environment, and thus under global change conditions, owing the timescale over which plastic phenotypic change occurs is short enough to reduce the time an organism spends in a novel environment with a suboptimal phenotype (Dupont et al., 2024; Padilla & Adolph, 1996).

### Acclimation environments drive the response to acute temperature changes

Beside the rate of plasticity, it is also critical to consider the temporal variations of environmental conditions. While the predictability of environmental changes has been largely explored in the study of plasticity evolution (Botero et al., 2015; Gavrilets & Scheiner, 1993; Leung et al., 2020; Reed et al., 2010; Tufto, 2015), the effect of acclimation environment on the level of plasticity has received relatively little attention. In this study, short-term temperature exposures suggested that the species performed better when acclimated at 5.0 °C rather than 7.5 °C, both in term of DETs number and functions, and reaction norm shape. Under the acute temperature exposure, fewer DETs were observed for fish acclimated at 5 °C compared to those at 7.5°C. In addition, a linear reaction norm and a bell or U-shape reaction norm were observed for fish acclimated at 5.0 and 7.5 °C, respectively. This result could be an indicator of the species’ performance in response to thermal stress. The bell shape response observed at 7.5 °C acclimation temperature is a pattern often observed in thermal performance curves in physiology. These curves are usually characterized by an initial slow increase performance with rising temperature, a maximum at some intermediate temperature, and then a rapid decrease at higher temperature (Angilletta, 2009; Huey & Stevenson, 1979). In our study, this pattern in the reaction norm of fish acclimated at 7.5 °C suggests a reduced capacity of *S. fasciatus* to cope with 2.5 °C temperature fluctuations. This hypothesis is also supported by the higher number of stress-related transcripts involved in the acute temperature response for fish acclimated at 7.5 °C compared to those at 5.0 °C expression which involved changes in growth-related transcripts.

Our results suggest that acclimation alone does not prepare species to cope with global warming, if considering the short-term transcriptional responses as a performance proxy. Our results showed that acclimation at 5.0 °C resulted in less stress-related response to fluctuating temperature compared to acclimation at 7.5 °C. Thus, our results fail to support the Beneficial Acclimation Hypothesis (BAH), in line with a significant number of empirical studies (Angilletta, 2009; Deere & Chown, 2006; Huey et al., 1999; Leroi et al., 1994; Wilson & Franklin, 2002). On the contrary, our results tend to show an Optimal developmental Temperature Hypothesis (OTH), and give support to a more specific one, namely the Cooler is Better Hypothesis (CBH). Under the CBH, organisms raised at cooler temperatures are expected to exhibit higher relative fitness across all temperatures compared to those raised at higher temperatures. This is based on the assumption that the advantages of large body size in ectotherms, developed at low temperature outweigh any benefits of acclimation (Huey et al., 1999; Wilson & Franklin, 2002).

Organisms following the CBH are more vulnerable to warning ocean. Consequently, our results show the limited ability of *S. fasciatus* to cope with temperature fluctuations through transcriptional plasticity under global change scenarios. In its natural habitat, *S. fasciatus* populations experience the range of temperature variation tested in this experiment due to the ongoing warming of the St. Lawrence System (Galbraith et al., 2024). Additionally, short-term temperature fluctuation (±2.5 °C) may occur due to their developmental or daily feeding vertical migrations (Gascon, 2003). As the temperature at which *S. fasciatus* lives in the St. Lawrence System increased from 5–6 °C to around 7.0 °C in the last 15 years (Lavoie et al., 2020), our results show that further increases in temperature, as expected with global warming, may surpass the species’ abilities to response to a warming environment through plasticity. Essentially, despite the capacity for thermal plasticity, an increase in ocean temperatures can significantly impact the initial phenotype of marine organisms. This change will potentially alter their level of plasticity, thereby affecting their ability to cope with environmental fluctuations. Instead, other mechanisms, like migration or other specific behavior, would be crucial for ectothermic organisms to cope with extreme temperatures (Gunderson & Stillman, 2015).

### Genetic variation in phenotypic plasticity

Within the St. Lawrence System, *S. fasciatus* exhibits notable genetic diversity. A recent population genomic study has identified at least three distinct populations, with instances of multiple populations being captured in a single bottom-trawl (Benestan et al., 2021). In our study, we conducted our experiments with animals captured in the St. Lawrence Estuary that were not studied in Benestan et al. (2021). Thus, we used ddRAD-seq genotypes from the *S. fasciatus* studied for transcriptomics and other *S. fasciatus* from the St. Lawrence System to understand if the individuals used in this study were associated to one or multiple populations (Supplementary Material 1). The specimens for this transcriptomic study formed a relatively genetically homogeneous group, distinct from other genetic groups of S*. fasciatus* observed in the St. Lawrence System (Fig. S1). The adaptive genetic variance may differ among these *S. fasciatus* genetic groups or populations. It is plausible that some populations have greater thermal tolerance than the *S. fasciatus* population we tested. Therefore, the potential for local adaptation to cope with increasing temperature is a valid consideration. For example, previous study of population genetics have identified a distinct population in the Gulf of Maine (Valentin et al., 2014), where individuals encounter areas with temperature reaching up to 13.0 °C (Scott, 1982).

Capacity for phenotypic plasticity has also a genetic basis which results in varying levels of plasticity among genetically diverse individuals and provides a source for natural selection to act upon, underscoring its evolutionary significance (Gavrilets & Scheiner, 1993; Landy et al., 2020; Leung et al., 2023; Scheiner, 1993). Recent evidence has also shown that rates of plasticity have diverged among different classes of ectotherms (Einum & Burton, 2023). This demonstrates that the rate of plasticity is also an evolving trait, highlighting its significant role in evolutionary ecology. Therefore, unraveling the genetic underpinnings of phenotypic plasticity will provide valuable insights into how species adapt to changing environment, and the potential strategies they may employ for survival. Genetic variation observed in *S. fasciatus* natural population could therefore underline variable capacity for and rate of plasticity.

### Gene functions involved in short- and long-term responses

Mechanisms underlying responses to temperature changes in stenothermal fish are not well understood (Ern et al., 2023). Our analysis of gene functions sheds light on the molecular response to fluctuating temperature for *S. fasciatus*, a temperate and stenotherm species. Short and long-term temperature exposures identified epigenetic, stress-related and heat coping molecular mechanisms associated to thermal responses. Our study also identified a small proportion of heat shock proteins (HSPs) and, surprisingly, genes related to eye functions underlying response to temperature changes in *S. fasciatus* muscles.

For both long and short-term responses to temperature change, GO analysis revealed rapid involvement of epigenetic processes, in particular those implicated in chromatin rearrangement via histone acetylation (Turner, 2000) and protein folding (Azzaz & Fantini, 2022; Harvey et al., 2018). Epigenetic mechanisms are defined as any chemical modifications that regulate gene expression pattern without any change in DNA sequence. The epigenome (all epigenetic modification of the genome of an organism) could be environmentally responsive and has thus been proposed as a major mechanism underlying phenotypic plasticity (Angers et al., 2010; Bollati & Baccarelli, 2010; Leung et al., 2023; Simola et al., 2016). The extent to which an organism can respond to current and future temperature extremes via epigenetic processes could influence its capacity to cope with global change (Bogan et al., 2020; Rey et al., 2020).

The stress-related functions involved in both short-term and long-term responses to temperature variation highlight a certain vulnerability of the species to increasing temperature. This is clearly demonstrated by the higher prevalence of stress-related functions in the short-term response, particularly in fish acclimated at the high temperature. After 24 h of exposure, response to temperature changes were mainly characterized by stress-related and energy production functions. Short-term temperature changes typically involve HSPs in fish (Jeffries et al., 2016; Liu et al., 2013; Tomanek, 2010). In this study, out of 125 transcripts identified as belonging to HSP gene family, only a few (N = 25) exhibited a heat shock response, while the majority were expressed at the same level across all experimental temperatures. The thermal niche of *S. fasciatus* is currently unknown, while could be inferred from population distribution in the field, but our results suggest it is narrow at least for this population from the St. Lawrence system. For example, HSP genes are found to be constantly expressed in Antarctic species to counteract the elevated levels of protein damage in cold environments (Clark, Fraser, Burns, et al., 2008; Place et al., 2004; Place & Hofmann, 2005). Moreover, variation in expression patterns and levels of induction could be observed among HSP70 gene isoforms, as described in the clam *Laternulla elliptica* (Clark, Fraser, & Peck, 2008). Our results also suggest the presence of other heat coping molecular mechanisms, akin to the function of HSPs. We observed that genes associated to cell growth and organization, protein folding or biogenesis were upregulated at higher temperature, whereas cell communication, transmembrane transport, and oxidoreductase activity genes were downregulated. These functions hint at role similar to those of HSPs, which are known to play a role in refolding damaged proteins (Feder & Hofmann, 1999).

We identified eye-related genes functions that responded to both long and short-term temperature changes in *S. fasciatus* muscles, potentially due to gene duplication and diversification of genes or a pleiotropic role of the same gene. The rhodopsin, an eye function gene, has been identified as important for adaptation to depth in multiple marine fish species. Specifically, positive selection based on depth habitat was detected repeatedly in multiple fishes, including *Sebastes* species (Larmuseau et al., 2011; Pampoulie et al., 2015; Schott et al., 2014; Sivasundar & Palumbi, 2010). Our results suggest that *S. fasciatus* genes annotated with eye-related functions are also being involved in temperature responses in the muscle tissue. Changes in the functions of vision-associated genes are well-documented across animal species. Zebrafish, for instance, exhibit a remarkable diversity in opsin genes, most of these genes being associated with non-visual functions such as regulation of the circadian rhythms, light-seeking behaviors, and seasonal responses (Davies et al., 2015). The opsin gene diversity likely emerged following genome duplications, and the retention of these genes is hypothesized to provide zebrafish with an adaptive advantage in bright environments. However, the functions of most of these non-visual opsin genes remain unconfirmed. Current research on opsin genes primarily targets eye tissues (e.g., Boyette et al., 2024) but our results suggest that these genes may instead, or also, be involved in sensing response to rapid temperature changes. For instance, the pleiotropic role of the *six6* genes, first identified as involved in the development of the eye and then important in the sensory ecology, has been demonstrated by identifying its expression in various tissues and developmental timepoints in Atlantic salmon (Moustakas-Verho et al., 2020).

## Conclusion

In conclusion, using transcriptomic analysis to assess the thermal plasticity of *S. fasciatus*, we shed light on molecular mechanisms underpinning plasticity. In addition, we emphasized the significant role of an organism’s past environment in determining its level of plasticity. Fish acclimated to higher temperatures exhibited an increased stress-related response to environmental fluctuations. This finding underscores the species’ susceptibility to ocean warming, despite its capacity for plasticity in response to changing temperature. Our study also highlights the value of examining the temporal dynamics of both environmental conditions and plastic response, alongside the capacity for plasticity. This approach enhances our understanding of the adaptiveness of plasticity and our ability to forecast eco-evolutionary responses to environmental changes.

## Acknowledgments

We thank D.C. Chavarria, J. Gagnon, D. Picard, T. Hansen, F. Hartog, K. MacGregor, JD. Tourangeau-Larivière and J. Heinerth for technical support during the experiment at the DFO Maurice Lamontagne Institute (Mont-Joli, QC, Canada) and for fish collection. We also acknowledge G. Bardaxoglou, J. Larivière and G. Cortial for DNA and RNA extraction.

## Data accessibility statement

Raw sequences data including the *S. fasciatus* genome assembly (Genbank accession: JBJQUQ000000000), the ddRAD-seq and RNA-seq datasets and metadata are available in the Sequences Read Archive (SRA) under the BioProject accession no. PRJNA1208449.

## Author contributions

DC, GP, CS and CL conceived the project and hypotheses. DC, CS and GP were responsible for funding acquisition. DC designed and supervised the tank experiment. JG ran the tank experiment. AB performed the bioinformatic analyzes of ddRADseq data. CL analyzed the transcriptomic data, prepared figures and tables, and wrote the original draft. All authors contributed to the article and approved the submitted version

## Supplementary materials

### <u>Supplementary material 1:</u> Population structure analysis

The DNA of 72 *Sebastes* from the basin experimental design (including the 48 used in the transcriptomic experiment) was extracted from muscle with DNeasy Blood and Tissus or DNeasy 96 (Qiagen) and quantified on a Synergy LX (BioTek) using PicoGreen to confirm high quality DNA extracts. Double digest restriction-site-associated DNA (ddRAD; *PstI* and *MspI* enzymes) libraries were prepared by the Plateforme d’Analyse Génomique (IBIS, Université Laval) using 20 ng of DNA per sample following Poland et al. (2012). Libraries were sequenced on a NovaSeq 6000 S4 150 PE at Genome Quebec, with 10% PhiX.

Genotyping was performed using STACKS modules (v.2.55, (Catchen et al. 2013; Rochette et al. 2019). Read quality was checked using FastQC (Andrews 2010) and multiQC (Ewels et al. 2016), and Illumina adaptors were removed with Trimmomatic (Bolger et al. 2014). Reads were demultiplexed with *process_radtag* module, with a truncation at 135 pb, then aligned to a *Sebastes fasciatus* genome (Genbank accession: JBJQUQ000000000) using BWA-MEM (Li and Durbin 2009) with default parameters.

We used the *gstacks* module to create two different SNP catalogs, 1) the species catalog to confirm that all individuals were *S. fasciatus* and 2) the population catalog to check for individual difference linked to population structure. To reach these objectives, we included in our analysis individuals from a fine scale population structure ddRAD project with similar DNA extraction and library preparation methods (Bourret et al. in prep). For the species catalog, we added 97 individuals representative of *S. fasciatus*, *S. mentella* and *S. norvegicus*. For the population catalog, we added 67 individuals presentative of the genetic diversity within *S. fasciatus*. From these two catalogs, different filtration steps were performed to keep only SNPs shared by > 75% individuals (*population* modules, r = 0.75, MAF 0.05), low missingness SNPs (<10%), to remove SNPs with observed heterozygosity (H_O_) > 0.6 and to keep only 1 SNP by loci. The resulting species and population dataset comprised 35,910 SNPs and 10,863 SNPs, respectively. Population structure was assessed using principal component analysis (PCA) using the *glPCA* function of adegenet R package (Jombart et al. 2008). All the individuals from the tank experiment, including the 48 transcriptomic ones, were confirmed to be *S. fasciatus* (Figure S1A), and most of them were genetically similar (Figure S1B).

**Fig. S1.**
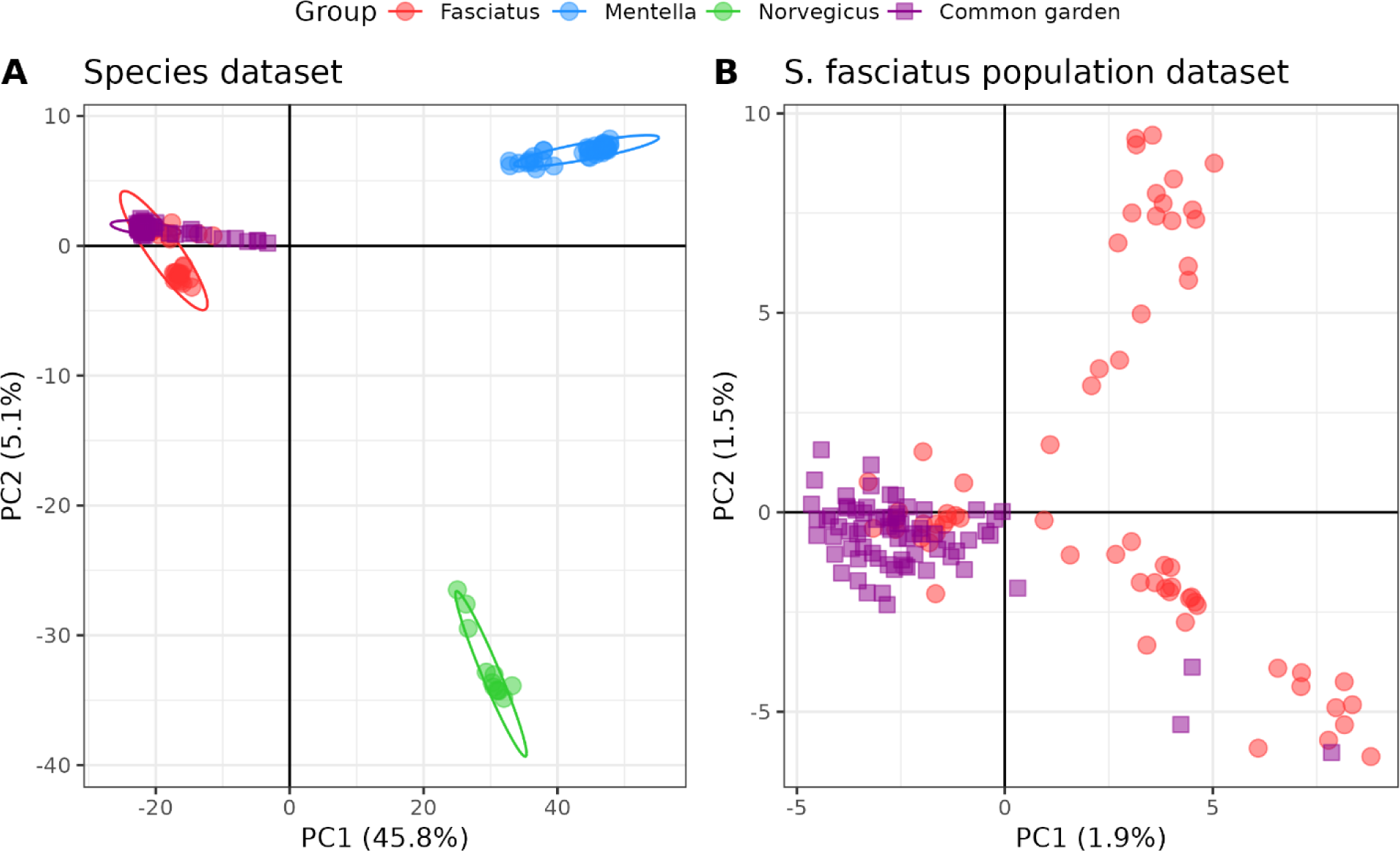
Genetic variation. Principal component analysis of genetic variation observed at the individual level for A) the species dataset and B) the population dataset. Common garden refers to the tank experiment.

**Fig. S2.**
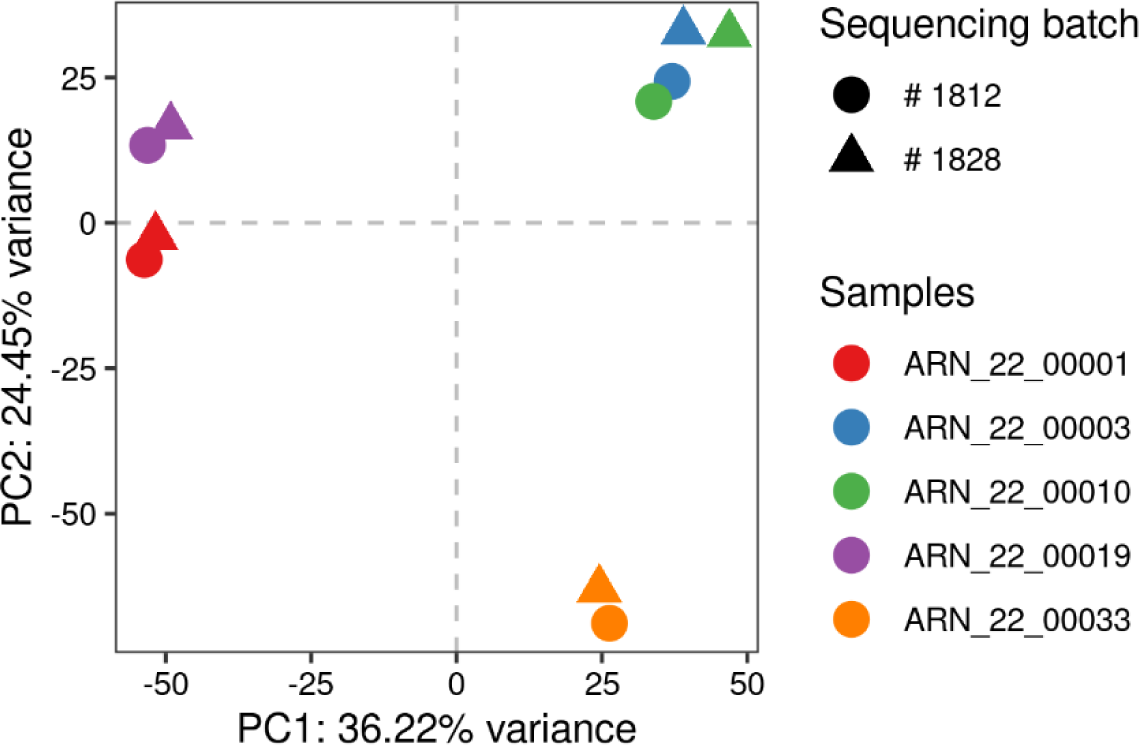
Sequencing batch effect on measured transcript expression level. Principal component analysis of transcript expression level for samples (colors) sequenced through different sequencing batches (shapes). Redundancy analysis revealed that sequencing batch as no effect on measured transcript expression level (*adjusted R^2^* = 3,66 %; P = 0.131). Only five up to the six samples sequenced in both sequencing batches are illustrated, given that one of them displayed an excess of rRNA (see Results section).

**Fig. S3.**
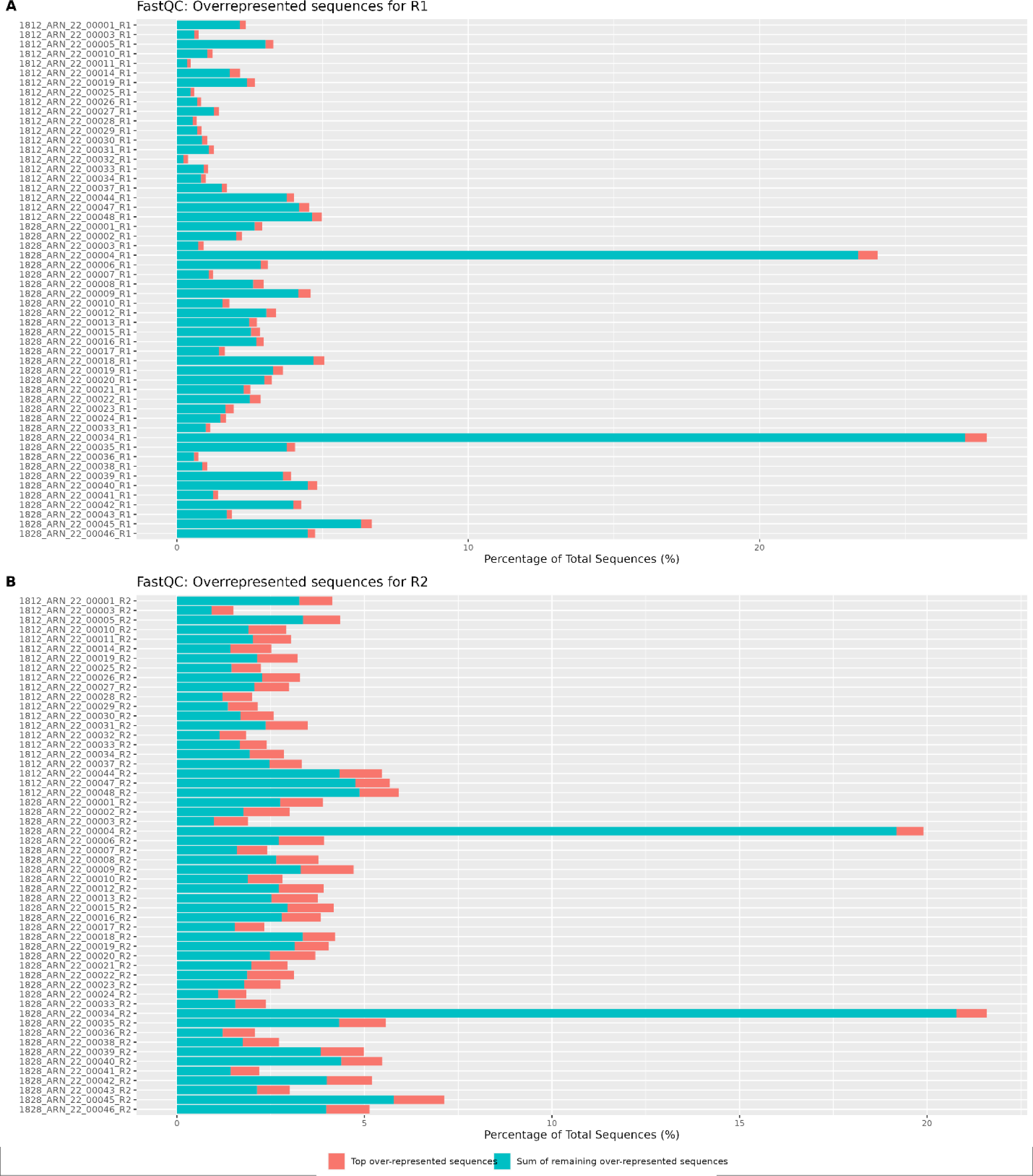
FastQC and MultiQC output of overrepresented sequences. R1 (A) and R2 (B) sequence quality check were performed on trimmed reads, after TrimGalore! Steps.

**Fig. S4.**
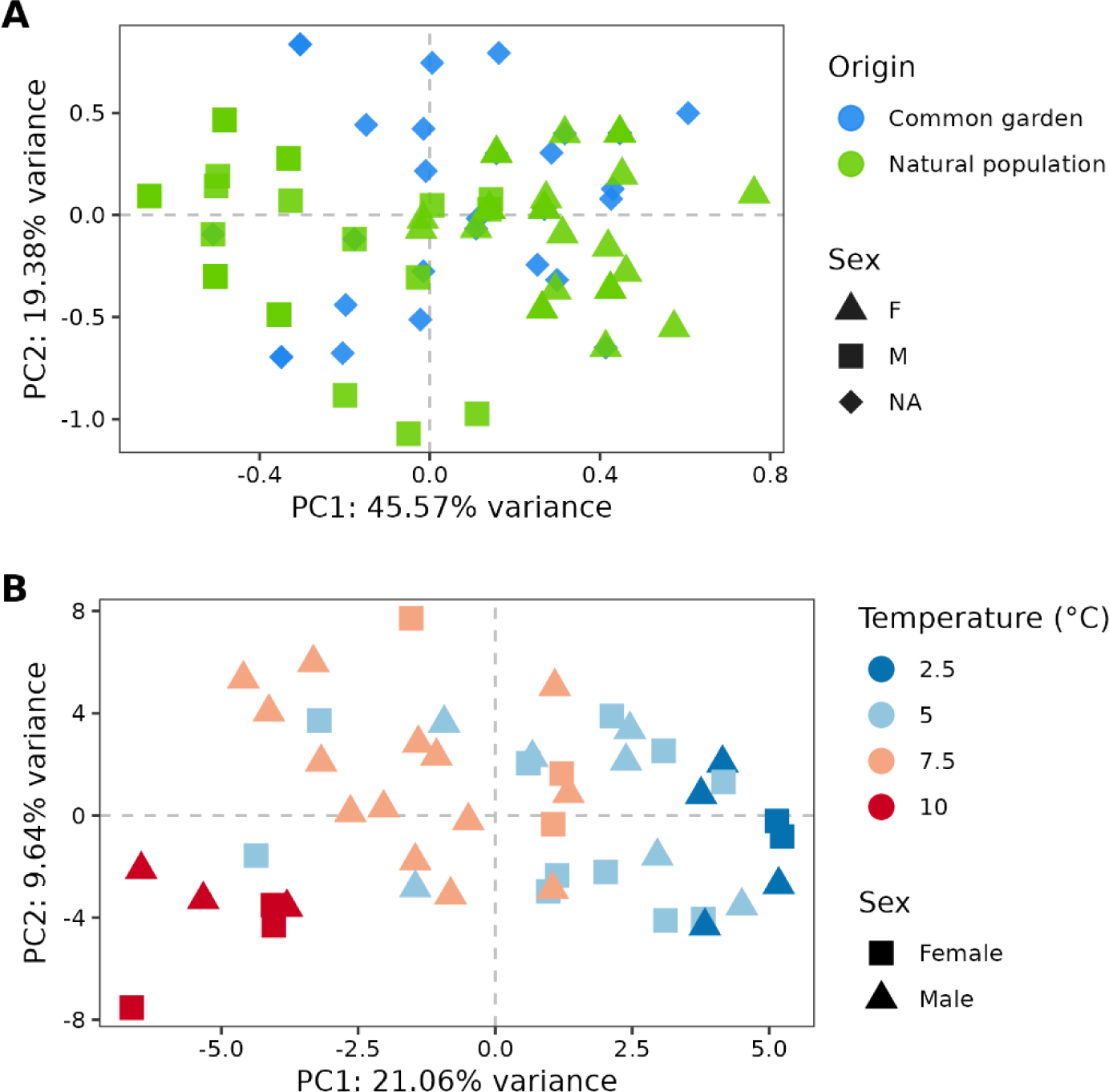
Sex effect on gene expression. **A. Genetic differentiation between male and female.** Principal component analysis (PCA) on six sex-specific SNPs for individuals from natural environments (green, Natural population) with known sex and used to identify sex (shape, square for female and triangle for male) of individuals from the basin experiment (blue, Common garden). Genetic data and samples from natural environments came from a distinct study using dd-RAD sequencing (Supp. Mat. 1). Briefly, sex visual identification were performed on adult and mature individuals (lengths > 220 mmm). AssignPOP (Chen et al. 2018) was used to identify sex-linked SNPs and to assign sex of common garden individuals. **B. Sex effect on gene expression.** PCA on transcript expression level of different sex (shape) acclimatized in four temperature (color), for individuals used for the RNA-seq analysis. Chen K-Y, Marschall EA, Sovic MG, Fries AC, Gibbs HL, Ludsin SA. assignPOP: An r package for population assignment using genetic, non-genetic, or integrated data in a machine-learning framework. Methods Ecol Evol. 2018; 9: 439–446. doi.org/10.1111/2041-210X.12897

**Fig. S5.**
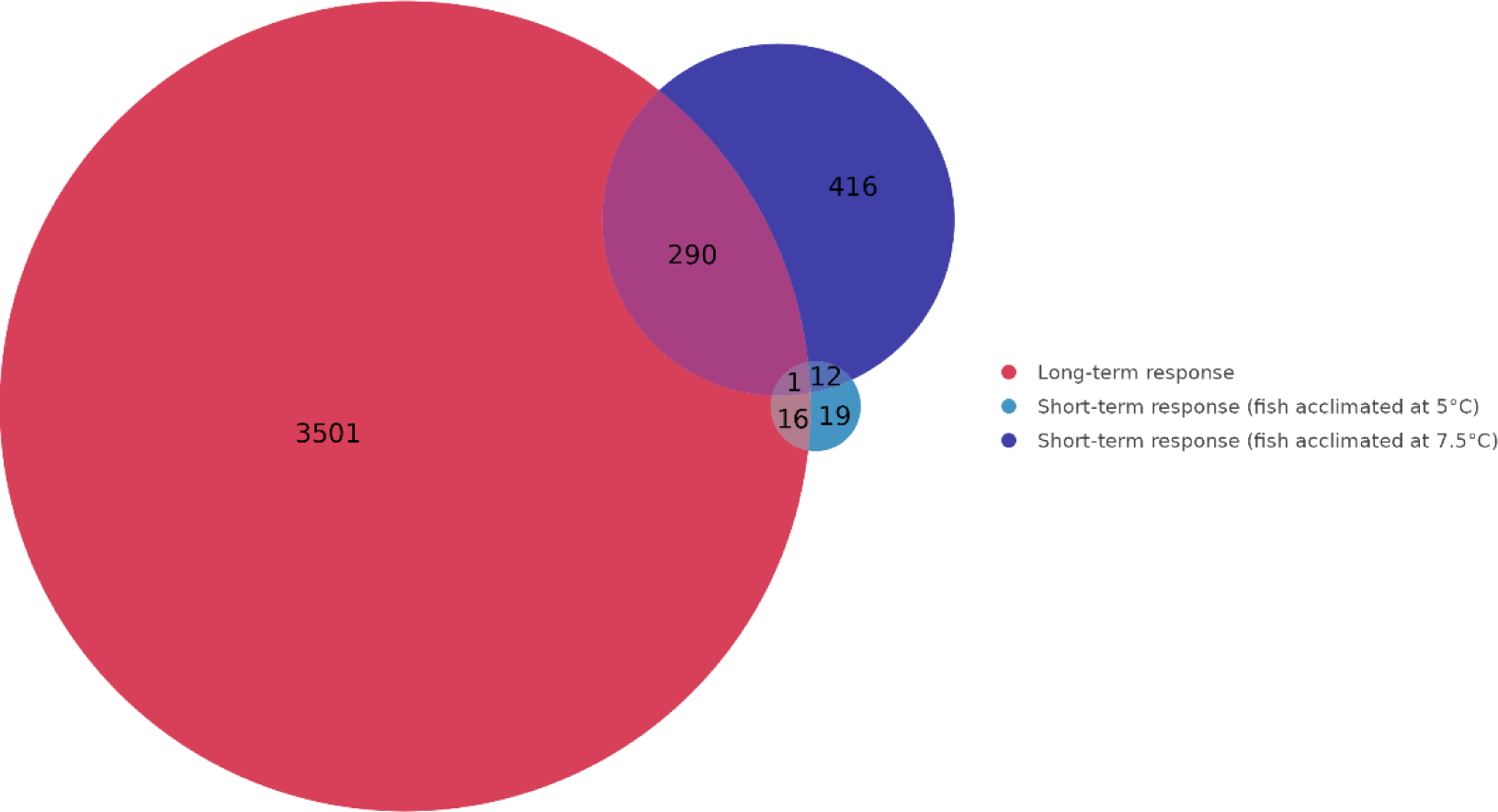
Differentially expressed transcripts (DETs) among all laboratory experiments. Venn diagram representing the number of DETs in response to long and short-term temperature stress exposure.

**Fig. S6.**
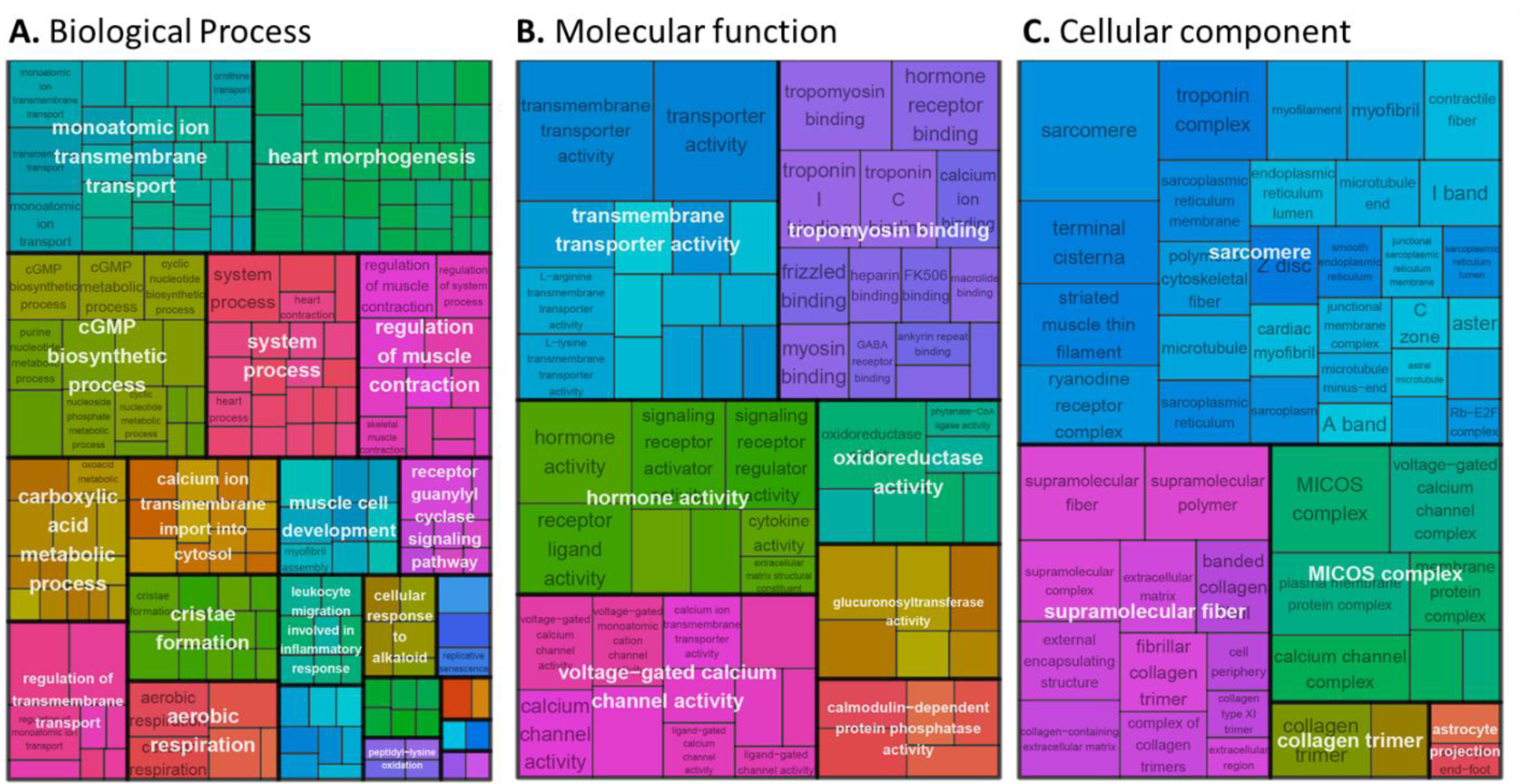
GO terms associated to group L01 of DETs in the long-term response to temperature change. Group L01 DETs is characterized by decreasing gene expression level when temperature increased (see Fig. 1C). Treemap for (A) biological process, (B) molecular function and (C) cellular component where GO terms were grouped (color) based on their semantic similarity, and the space used by the term is proportional to the −log10(adjusted *P-value*), hence the gene function candidate probability.

**Fig. S7.**
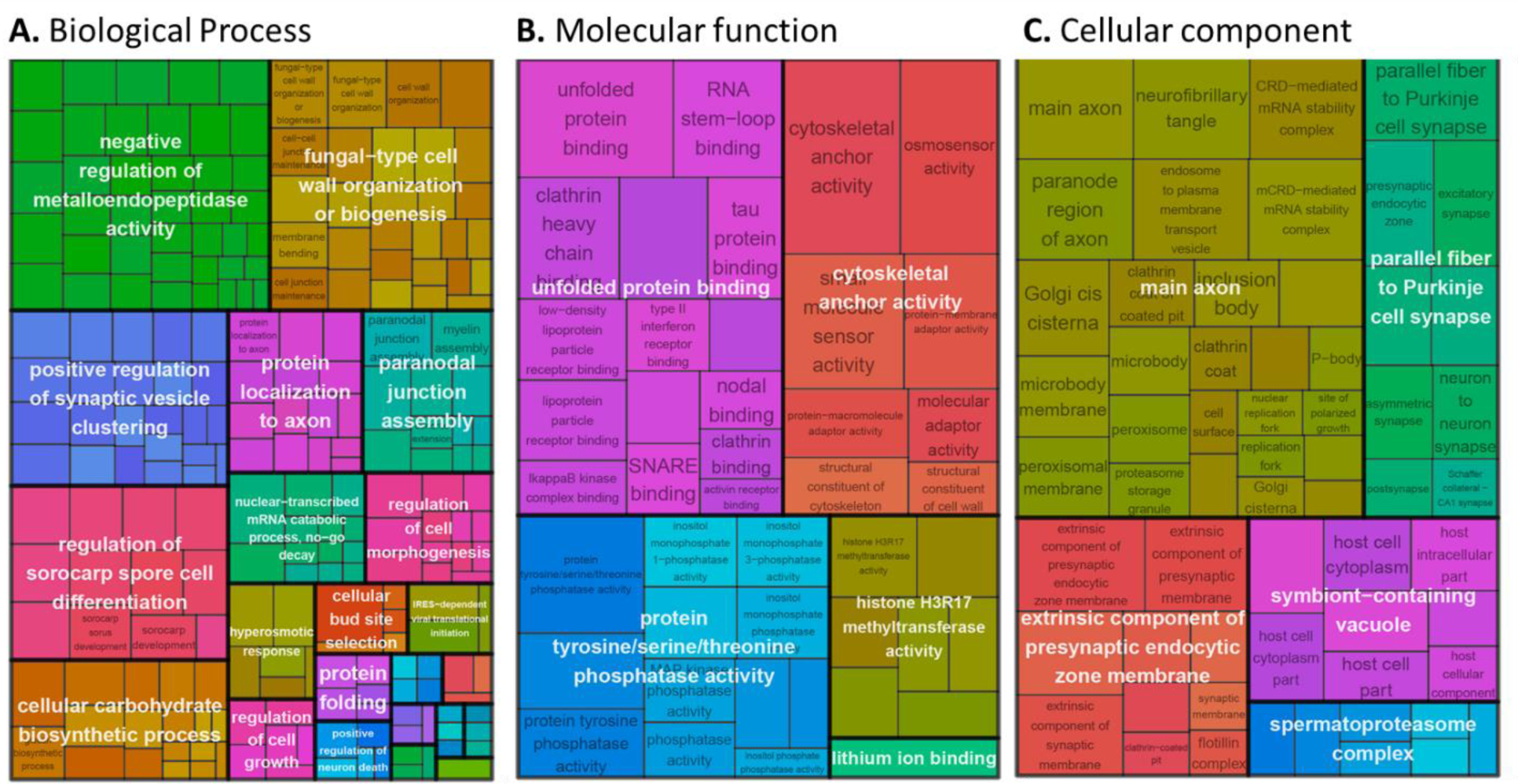
GO terms associated to group L02 of DETs in the long-term response to temperature change. Group L02 DETs is characterized by increasing gene expression level when temperature increased (see Fig. 1C). Treemap for (A) biological process, (B) molecular function and (C) cellular component where GO terms were grouped (color) based on their semantic similarity, and the space used by the term is proportional to the −log10(adjusted *P-value*), hence the gene function candidate probability.

**Fig. S8.**
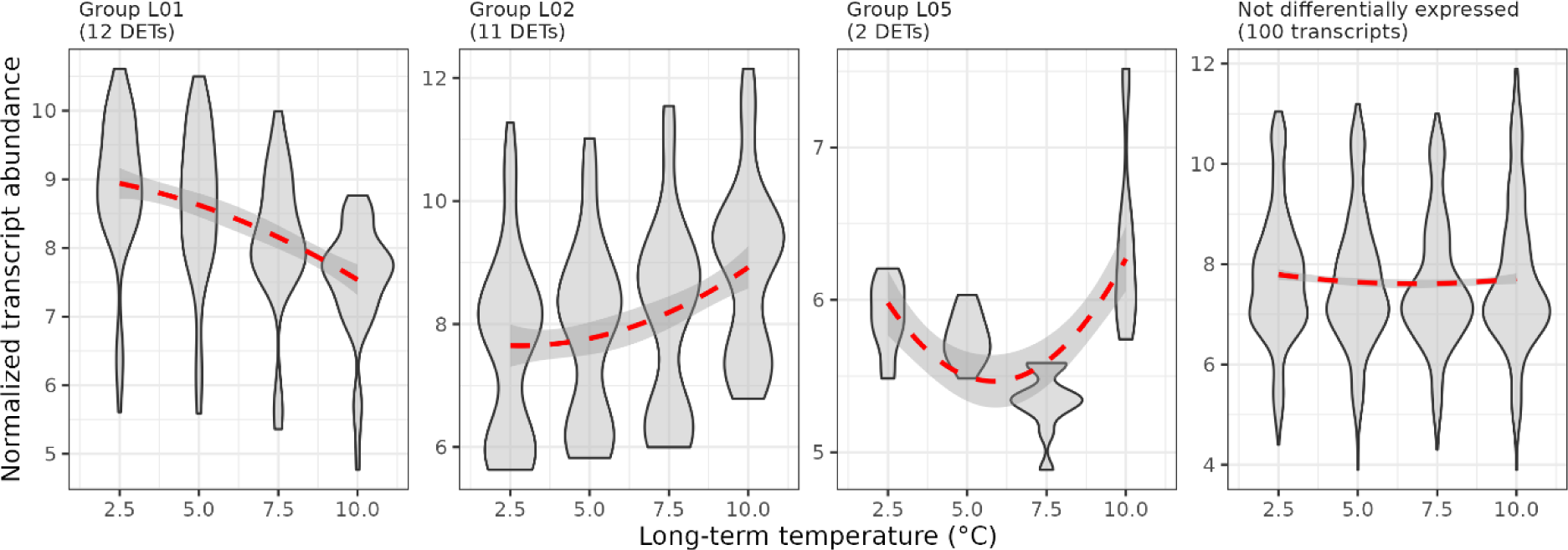
Expression level of transcript associated to heat-shock proteins (HSPs). Normalized transcript abundance were obtained using the variance stabilizing transformation as implemented in DESeq2. Transcripts associated to HSPs and detected to be differentially expressed (DETs) among temperatures were grouped according to the expression moduled they belong to, including transcripts that were down reglulated (Group L01) or upregulated at higher temperatures (Group L02 and L05).

